# Light-dependent induction of *Edn2* expression and attenuation of retinal pathology by endothelin receptor antagonists in *Prominin-1*- deficient mice

**DOI:** 10.1101/2020.12.07.405308

**Authors:** Yuka Kobayashi, Shizuka Watanabe, Manabu Shirai, Chiemi Yamashiro, Tadahiko Ogata, Fumiaki Higashijima, Takuya Yoshimoto, Takahide Hayano, Yoshiyuki Asai, Noriaki Sasai, Kazuhiro Kimura

## Abstract

Retinitis pigmentosa (RP) and macular dystrophy (MD) are prevalent retinal degenerative diseases associated with gradual photoreceptor death. These diseases are often caused by genetic mutations that result in degeneration of the retina postnatally after it has fully developed. The *Prominin-1* gene (*Prom1*) is a causative gene for RP and MD, and *Prom1*- knockout (KO) mice recapitulate key features of these diseases including light-dependent retinal degeneration and stenosis of retinal blood vessels. The mechanisms underlying progression of such degeneration have remained unknown, however. We here analysed early events associated with retinal degeneration in *Prom1*-KO mice. We found that photoreceptor cell death and glial cell activation occur between 2 and 3 weeks after birth. High-throughput analysis revealed that expression of the endothelin-2 gene (*Edn2*) was markedly up-regulated in the Prom1-deficient retina during this period. Expression of *Edn2* was also induced by light stimulation in *Prom1*-KO mice that had been reared in the dark. Finally, treatment with endothelin receptor antagonists attenuated photoreceptor cell death, gliosis, and retinal vessel stenosis in *Prom1*-KO mice. Our findings suggest that inhibitors of endothelin signalling may delay the progression of RP and MD and therefore warrant further study as potential therapeutic agents for these diseases.

## 1. Introduction

Both retinitis pigmentosa (RP) and macular dystrophy (MD) are inherited retinal disorders associated with progressive photoreceptor cell death [1]. These diseases have a combined prevalence of 1 in 3000 to 4000 people worldwide. Initial symptoms include nyctalopia (night blindness) and visual field deficits, which are followed by loss of visual acuity and colour blindness and eventually by complete blindness. Although >60 genes encoding various types of protein - including membrane proteins, transcription factors, splicing regulators, and enzymes related to the visual cycle - have been implicated in RP and MD [1], these conditions remain incurable, with effective therapeutic strategies remaining to be established, and they have profound effects on the quality of life.

The *Prominin-1* gene (*Prom1*, also known as *AC133*, *CD133*, and *RP41*) encodes a pentaspan transmembrane glycoprotein that is expressed in photoreceptor cells of the retina as well as in kidney and testis [2]. Several mutations of *Prom1* have been identified in individuals with RP or MD [3–5], with all such mutations resulting in amino acid substitutions or carboxyl-terminal truncations of the encoded protein. The mechanisms underlying RP and MD associated with *Prom1* mutations have been investigated by studies of several lines of *Prom1*-knockout (KO) mice [5–7]. Although photoreceptor cells develop normally in these KO mice, they begin to degenerate after birth, resulting in a progressive loss of the outer nuclear layer (ONL) of the retina and recapitulation of the signs of RP and MD. The retinal vasculature also becomes attenuated with disease progression [7].

We previously showed that photoreceptor cells of the *Prom1*-KO mouse retina degenerate in response to light stimulation. Such mice reared in a completely dark setting thus manifested a marked delay in the loss of photoreceptor cells. We therefore suggested that the mutant retinal cells are hypersensitive to light stimulation and experience phototoxicity [6]. The visual cycle was also found to be impaired in the *Prom1*-KO cells, and treatment based on chemical compounds that modulate the visual cycle was found to mitigate the mutant phenotype [6].

The Prom1 protein localises to the connecting cilium and outer segment of both rod and cone photoreceptors [3]. Ultrastructural analysis revealed the structure of the outer segment to be severely disorganised in photoreceptor cells of *Prom1*-KO mice, whereas other photoreceptor components - including the inner segment, nucleus, and axon - remained largely intact [6, 7]. Biochemical analysis has shown that two tyrosine residues in the carboxyl-terminal region of Prom1 are phosphorylated by the tyrosine kinases Src and Fyn, although the physiological implications of such phosphorylation remain to be elucidated [8]. Prom1 has also been shown to interact with the p85 regulatory subunit of phosphatidylinositol 3-kinase (PI3K) and to be essential for both the self-renewal and tumourigenic capacity of glioma stem cells [9]. In addition, Prom1 has been detected in cilia, which are protrusive structures at the cell membrane and key signalling hubs [10], and to be essential for maximisation of Hedgehog signalling in neural stem cells [11]. We recently showed that Prom1 activates the small GTPase Rho and regulates chloride conductance triggered by intracellular calcium uptake [12].

To characterise the mechanisms underlying the role of Prom1 dysfunction in retinal degeneration and thereby to provide insight into potential treatments for *Prom1* mutation-associated RP and MD, we here investigated the initial manifestations of such degeneration. We analysed *Prom1* expression as well as the ONL transition in *Prom1*-KO mice. We then performed a high-throughput expression analysis to identify genes responsible for degeneration of the *Prom1*-deficient retina. Our results implicated an inflammatory pathway dependent on the endothelin 2 gene (*Edn2*), and we found that a chemical treatment targeted to endothelin signalling mitigated the deterioration of retinal structure and function in *Prom1*-KO mice.

## 2. Methods

### 2.1. Mice

*Prom1*-KO mice were established previously (CDB0623K, http://www2.clst.riken.jp/arg/methods.html), and they were reared on a hybrid genetic background of C57BL/6 and CBA/NSlc strains. The targeting vector for *Prom1* ablation contained the *lacZ* (β-galactosidase) gene, with the result that expression of this latter gene reflects that of *Prom1*. Both the *Prom1*-KO mice and their wild-type (WT) littermates were kept on a 12-hour-light, 12-hour-dark cycle, with the cage racks being covered with blackout curtains and all procedures including feeding and cage maintenance being performed in the absence of light (<0.5 lux) during the dark phase. For experiments involving light stimulation, mice were exposed for 3 h to a light panel (LED viewer 5000; Shinko, Tokyo, Japan) placed on top of the cage, which resulted in a light intensity of 3800 lux at the bottom of the cage. For chemical treatment, mice received intraperitoneal injections (2 mg/kg) of each of the endothelin receptor antagonists BQ-123 (ab141005, Abcam) and BQ-788 (ab144504, Abcam) on postnatal day (P) 14, P19, and P24. The mice were then subjected to analysis on P28.

### 2.2. RNA extraction and RT-qPCR analysis

The retina, retinal pigment epithelium (RPE), and testis were dissected from mice killed by cervical dislocation. Total RNA was extracted from the isolated tissue and was subjected to reverse transcription (RT) with the use of a NucleoSpin RNA extraction kit (U955C, Takara) and PrimeScript RT reagent kit (RR037, Takara), respectively. The resulting cDNA was subjected to quantitative polymerase chain reaction (qPCR) analysis with a CFX qPCR machine (Bio-Rad) and with primers listed in supplementary table S1. The amplification data were analysed with the comparative *C*_t_ method, and gene expression levels were normalised by that of the glyceraldehyde-3-phosphate dehydrogenase gene (*Gapdh*).

### 2.3. High-throughput expression analysis

Total RNA samples were prepared from three (P14) or four (P21) retinas of WT or *Prom1*-KO mice and were used to synthesise cDNA libraries with a TruSeq stranded-mRNA library preparation kit (Illumina, 20020594). The libraries were sequenced with the NextSeq 500 platform (Illumina). In total, approximately twenty million reads/sample were mapped with the CLC genomics workbench software (Qiagen) [13]. The sequencing data were deposited in the DNA Data Bank of Japan (DDBJ) public database, with the accession number of SSUB016168. Gene ontology (GO) term analysis was performed according to the Kyoto Encyclopaedia of Genes and Genomes database (KEGG, https://www.genome.jp/kegg).

### 2.4. Immunofluorescence analysis, β-galactosidase and isolectin staining, and TUNEL analysis

For immunofluorescence analysis, the enucleated retina was fixed for 2 h with a mixture of 1% paraformaldehyde and 0.2% glutaraldehyde in phosphate-buffered saline (PBS), incubated overnight in PBS containing 15% sucrose, embedded in O.C.T. compound (Sakura), and sectioned at a thickness of 12 μm. The sections were exposed to mouse monoclonal antibodies to GFAP (G3893; Sigma) or rabbit polyclonal antibodies to Iba-1 (019-19741; Wako), and immune complexes were detected with Cy3-conjugated secondary antibodies (715-166-151 and 715-166-152 for mouse and rabbit, respectively; Jackson Immunoresearch). Nuclei were counterstained with 4’,6-diamidino-2-phenylindole (DAPI) with the use of DAPI Fluoromount-G (0100-20; Southern Biotech). Sections were also stained for β-galactosidase (β-gal) activity with the use of a staining kit (11828673001, Roche). Apoptotic cells were detected by TUNEL analysis with digoxigenin-labelled dUTP (S7105, Merck Millipore), terminal deoxynucleotidyl transferase (3333566001, Merck), and rhodamine-conjugated antibodies to digoxigenin (11207750910, Roche). For preparation of flat-mount samples, the retina was fixed for 150 min with 4% paraformaldehyde and the RPE was peeled off. The samples were subjected to isolectin staining by consecutive exposure to 5% dried skim milk and Alexa Flour 488-conjugated GS-IB4 (I21411, Thermo Fisher Scientific) as described previously [14]. Images were acquired with an LSM 710 confocal microscope (Zeiss) for immunofluorescence, β-gal, and TUNEL staining, or with a BZ-X710 microscope (Keyence) for flat-mount preparations. Imaging data were processed and integrated with Photoshop (Adobe) and Illustrator (Adobe) software, respectively.

### 2.5. Statistical analysis

Quantitative data are presented as means ± s.e.m. Differences between two or among more than two groups were evaluated with the two-tailed Student’s *t* test and by one-way analysis of variance (ANOVA) followed by Tukey’s post hoc test, respectively. Statistical analysis was performed with Prism software (Graphpad), and a *p* value of <0.05 was considered statistically significant.

## 3. Results

### 3.1. *Prom1* is expressed in the retina from perinatal to adult stages

We previously showed that retinal cells in *Prom1*-KO mice appear to develop normally before the onset of degeneration [6]. We here first examined the spatiotemporal expression of *Prom1* in the mouse retina. Given that our *Prom1*-KO mice harbour the *lacZ* gene at the *Prom1* locus, we performed staining for β-gal activity in the heterozygous mutant mice at birth as well as at P2 (figure 1*a*-*a”*), P14 (figure 1*b*-*b”*), P21 (figure 1*c*-*c”*), and P42 (figure 1*d*-*d”*). At all the stages analysed, β-gal staining was localised predominantly to the outer layers in the retina, with more sporadic staining also apparent in the inner nuclear layer (INL). Given that retinal phenotypes of *Prom1*-KO mice are not obvious until 2 weeks after birth, these results suggested that *Prom1* expression precedes the onset of function of the encoded protein in postnatal retinal homeostasis.

**Figure 1.**
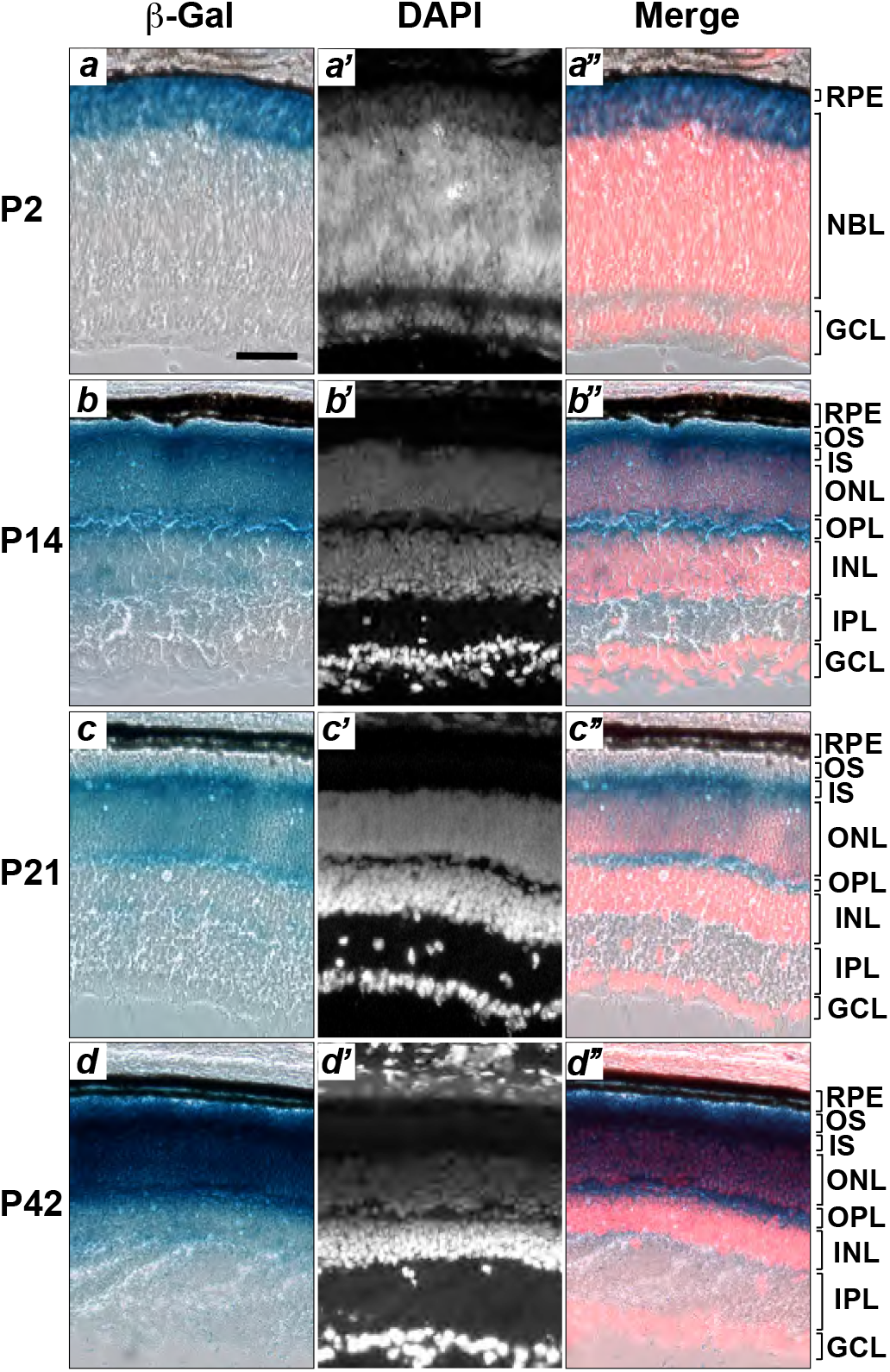
*Prom1* is expressed in the ONL of the retina from perinatal to adult stages. The retina of heterozygous *Prom1* mutant mice at P2 (*a-a”*), P14, (*b-b”*), P21 (*c-c”*), and P42 (*d-d”*) was subjected to staining of β-gal activity (*a*,*b*,*c*,*d*) as well as to staining of nuclei with DAPI (*a’*,*b’*,*c’*,*d’*). Merged images are also shown (*a”*,*b”*,*c”*,*d”*). Data are representative of three retinas at each age. Scale bar in (*a*) is (50 μm) and applies to all images. RPE, retinal pigment epithelium; NBL, neuroblast layer; GCL, ganglion cell layer; OS, outer segments; IS, inner segments; ONL, outer nuclear layer; OPL, outer plexiform layer; INL, inner nuclear layer; IPL, inner plexiform layer.

### 3.2. The *Prom1*-KO mouse retina manifests both apoptosis and an inflammatory response at 3 weeks after birth

We previously showed that the retina of *Prom1*-KO mice appears normal at P14 and begins to degenerate soon after the animals first open their eyes at P14 [6]. We therefore investigated whether the *Prom1*-deficient retina might undergo apoptosis in response to light exposure. Whereas the TUNEL assay revealed few apoptotic cells in the retina of WT or *Prom1*-KO mice at P14 (figure 2*a* and *b*), a significant increase in the number of TUNEL-positive cells, located mainly in the ONL, was detected at P21 in the *Prom1*-KO retina (figure 2*c*–*e*). These results suggested that programmed cell death by apoptosis begins to occur in the ONL of the retina between 2 and 3 weeks after birth in *Prom1*-KO mice.

**Figure 2.**
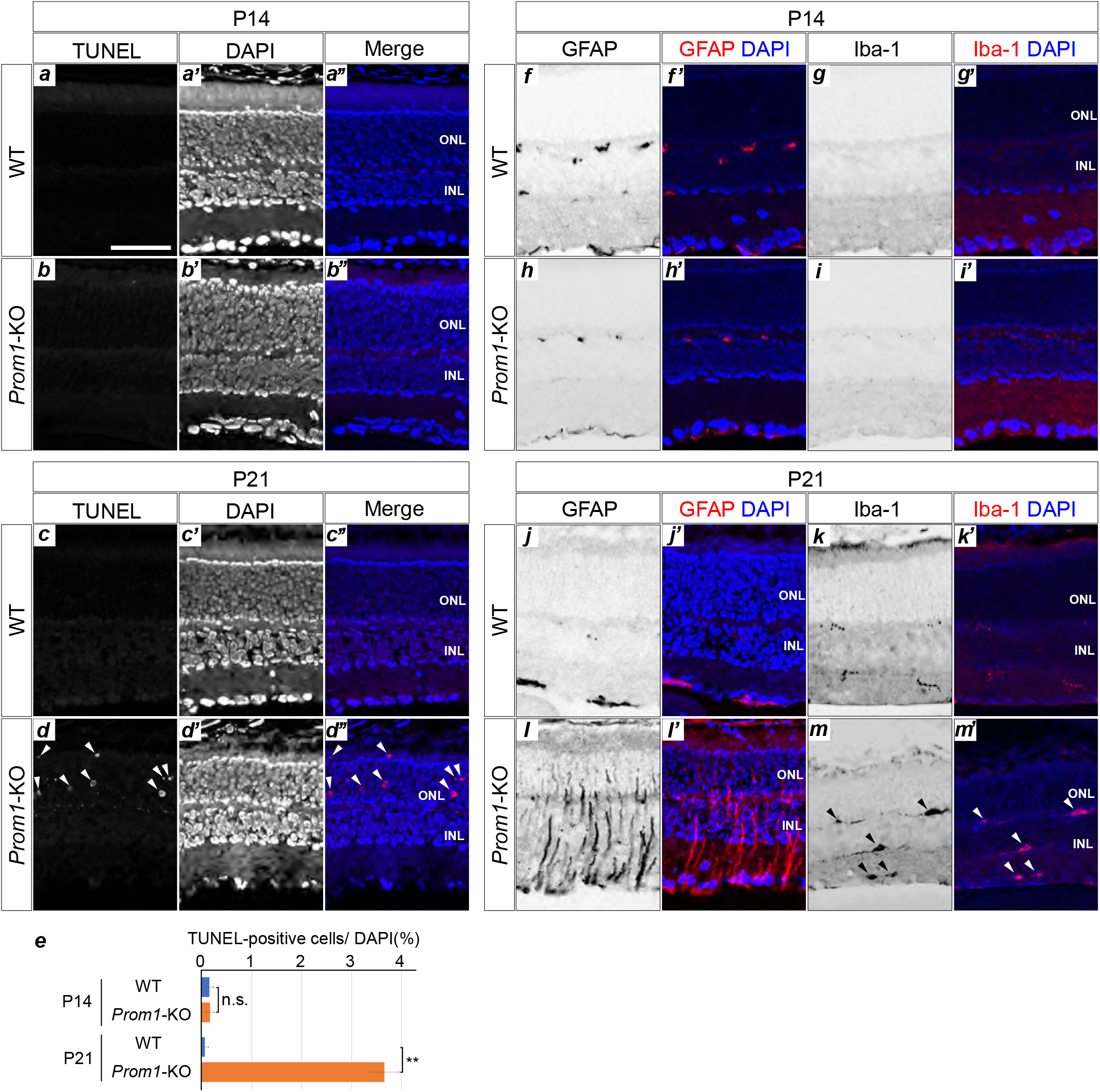
Programmed cell death and an inflammatory response in the postnatal *Prom1*-KO mouse retina. (*a*–*d”*) TUNEL staining of the WT (*a*-*a”*,*c*-*c”*) and *Prom1*-KO (*b*-*b”*,*d*-*d”*) mouse retina at P14 (*a*-*b”*) and P21 (*c*-*d”*). Nuclei were stained with DAPI (*a’,b’*,*c’*,*d’*). Merged images of TUNEL (red) and DAPI (blue) staining are also shown (*a”*,*b”*,*c”*,*d”*). Arrowheads in (*d*,*d”*) indicate apoptotic cells. (*e*) Quantitation of the proportion of TUNEL-positive cells among all DAPI-stained cells for images similar to those in (*a*),(*b*),(*c*) and (*d*). Data are means ± s.e.m. for four retinas for each condition. ** *p* < 0.01; n.s., not significant (two-tailed Student’s *t* test). (*f*–*m*) Immunofluorescence staining for GFAP (*f*, *h*,*j*,*l*) and Iba-1 (*g*,*i*,*k*,*m*) in the retina of WT (*f*,*g*,*j*,*k*) and *Prom1*-KO (*h*,*i*,*l*,*m*) mice at P14 (*f*–*i*) and P21 (*j*–*m*). Merged images with DAPI staining are also shown (*f’*,*g’*,*h’*,*i’*,*j’*,*k’*,*l’*,*m’*). Arrowheads in (*m*) indicate Iba-1– positive cells. Data are representative of three (P14) or five (P21) retinas for each genotype. Scale bar in (*a*) is 50 μm and applies to all images.

Glial fibrillary acidic protein (GFAP) is an intermediate filament protein that is expressed by Müller glia in response to retinal injury [15, 16]. Similarly, Iba-1 is a scaffold protein that is expressed in microglia and which is up-regulated during an inflammatory response [17, 18]. We therefore next examined whether the *Prom1*-KO retina might undergo light-induced inflammation by analysing the expression of these two proteins. Immunofluorescence analysis revealed that, whereas both GFAP and Iba-1 were essentially undetectable in the WT or *Prom1*-KO retina at P14 (figure 2*f*–*i*), a marked increase in the extent of staining for both proteins was observed in the *Prom1*-KO retina at P21 (figure 2*j*–*m*), suggesting that the increased cell death that occurs in the ONL of the mutant mice after birth is accompanied by the activation of glial cells.

### 3.3. Inflammation-related gene expression is up-regulated in the *Prom1*-KO mouse retina

We next sought to identify genes whose expression might be affected by Prom1 deficiency by subjecting the retina of WT and *Prom1*-KO mice at P14 and P21 to high-throughput expression analysis based on RNA sequencing. Gene expression at P14 tended to vary within each genotype, and the only gene whose expression differed significantly between genotypes was *Prom1* itself (figure 3*a*, supplementary table S2), suggesting that Prom1 does not significantly influence the gene expression profile at P14. In contrast, the expression of various genes differed between the two genotypes at P21 (figure 3*b*, supplementary table S3). The expression of 1,081 and 766 genes was thus up- and down- regulated, respectively, in the *Prom1*-KO retina with a *p* value of <0.01. In particular, expression of *Edn2* was the most consistently and markedly up-regulated in the *Prom1*-KO retina. The expression of genes associated with the inflammatory response - such as *Ifi44l*, *Serpina3n*, *S100a6*, *Bcl3*, and *Gfap* - was also increased in the *Prom1*-KO retina at P21. Conversely, the expression of genes related to RP or of those essential for retinal development and functional homeostasis - including *Fscn2* (RP30) [19], *Prph2* (RP7) [20], *Nr2e3* (RP37) [21], *Kcnv2* [22], *Elovl2* [23], *Pde6b* (RD1) [24], and *Ttc21b* [25] - was down-regulated in the *Prom1*-KO retina at P21 (supplementary table S3). GO term analysis revealed that several signalling pathways, including apoptotic (TNF) and infectious-related signal (Epstein-Barr virus infection) signals, were affected by the loss of Prom1 (figure 3*c*).

**Figure 3.**
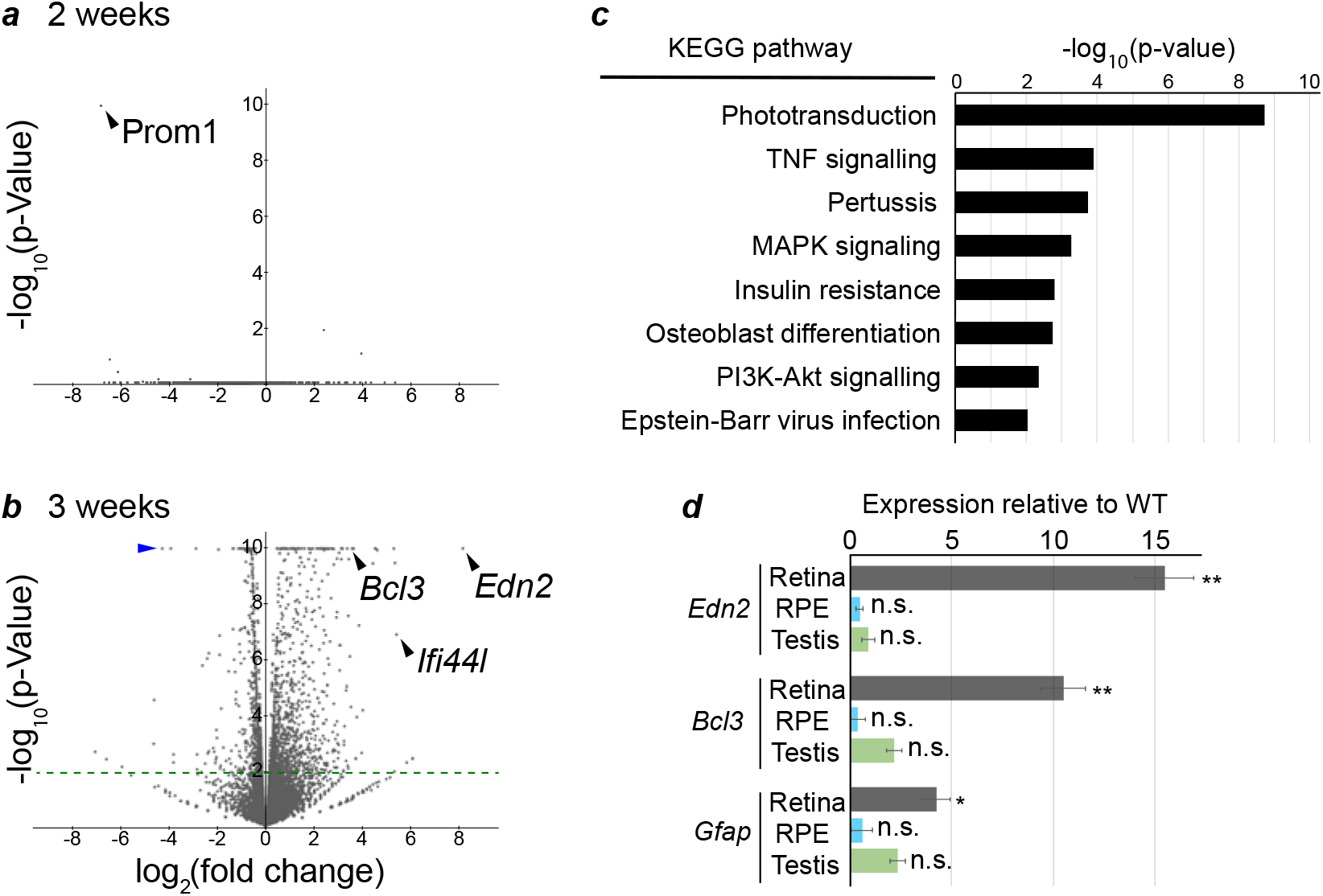
Effects of Prom1 deficiency on gene expression in the retina. (*a*,*b*) Volcano plots for RNA-sequencing analysis of the retina of *Prom1*-KO mice relative to that of WT mice at P14 (*a*) and P21 (*b*). Genes with a *p* value of 1 × 10^−10^ are indicated with the blue arrowhead in (*b*). A cut-off *p* value of 1 × 10^−2^ is indicated by the green dashed line. Data are for three (P14) or four (P21) retinas of each genotype. (*c*) GO term analysis based on KEGG pathways for genes whose expression differed significantly between the retinas of *Prom1*-KO and WT mice in the RNA-sequencing analysis at P21. (*d*) RT-qPCR analysis of *Edn2*, *Bcl3*, and *Gfap* expression in the retina, RPE, and testis of WT and *Prom1*-KO mice at P21. Data are means ± s.e.m. for three retinas of each genotype. \**p* < 0.05, \*\**p* < 0.01, n.s., not significant (two-tailed Student’s *t* test).

We also investigated whether the observed effects of Prom1 deficiency on gene expression were specific to the retina. Given that *Prom1* is expressed in the retina, RPE, and testis [2], we performed RT-qPCR analysis of RNA prepared from these tissues of WT and *Prom1*-KO mice at P21. Consistent, with the results of our RNA-sequencing analysis, the expression of *Edn2*, *Bcl3*, and *Gfap* was increased in the retina of *Prom1*-KO mice (figure 3*d*). However, the expression of these genes in the RPE and testis did not differ between the two genotypes, indicating that the effect of Prom1 on their expression is specific to the retina. Together, these various data suggested that Prom1 deficiency results in up-regulation of inflammation-related genes and down-regulation of genes essential for functional homeostasis of photoreceptor cells at 3 weeks after birth.

### 3.4. Inflammation-related gene expression is increased by light stimulation in the *Prom1*-KO mouse retina

To determine the mechanism underlying the up-regulation of specific gene expression apparent in the retina of *Prom1*-KO mice at P21, we examined whether light stimulation might play a role. We therefore compared such gene expression between P21 retinas obtained from *Prom1*-KO mice reared under a normal day-night cycle or in the dark. RT-qPCR analysis revealed that, whereas the expression of *Edn2*, *Bcl3*, and *Gfap* did not differ between *Prom1*-KO and WT mice reared in the dark condition, marked up-regulation of the expression of each of these genes was apparent specifically in *Prom1*-KO mice raised under the normal day-night condition (figure 4*a*). Consistent with these results, immunofluorescence analysis showed that the number of GFAP-positive cells in the retina was smaller for *Prom1*-KO mice reared in the dark compared with those reared under the normal condition (figure 4*b* and *c*). To examine further the effect of light on gene expression, we maintained *Prom1*-KO mice and their WT littermates under the dark condition for 3 weeks, exposed them to a bright light for 3 h, and then allowed them to recover for 3 days in the dark. The retina was then dissected and subjected to RT-qPCR and immunofluorescence analyses. Light stimulation resulted in a marked increase both in the expression of *Edn2* and *Bcl3* (Figure 4*d*) and in the number of GFAP-positive cells (figure 4*e*) in the retina of *Prom1*-KO mice but not in that of WT mice. Collectively, these results thus suggested that the up-regulation of *Edn2*, *Bcl3*, and *Gfap* expression apparent in the retina of *Prom1*-KO mice is an immediate response to light stimulation, and that the inflammatory response mediated by these genes is one of the primary events leading to degeneration of the mutant retina.

**Figure 4.**
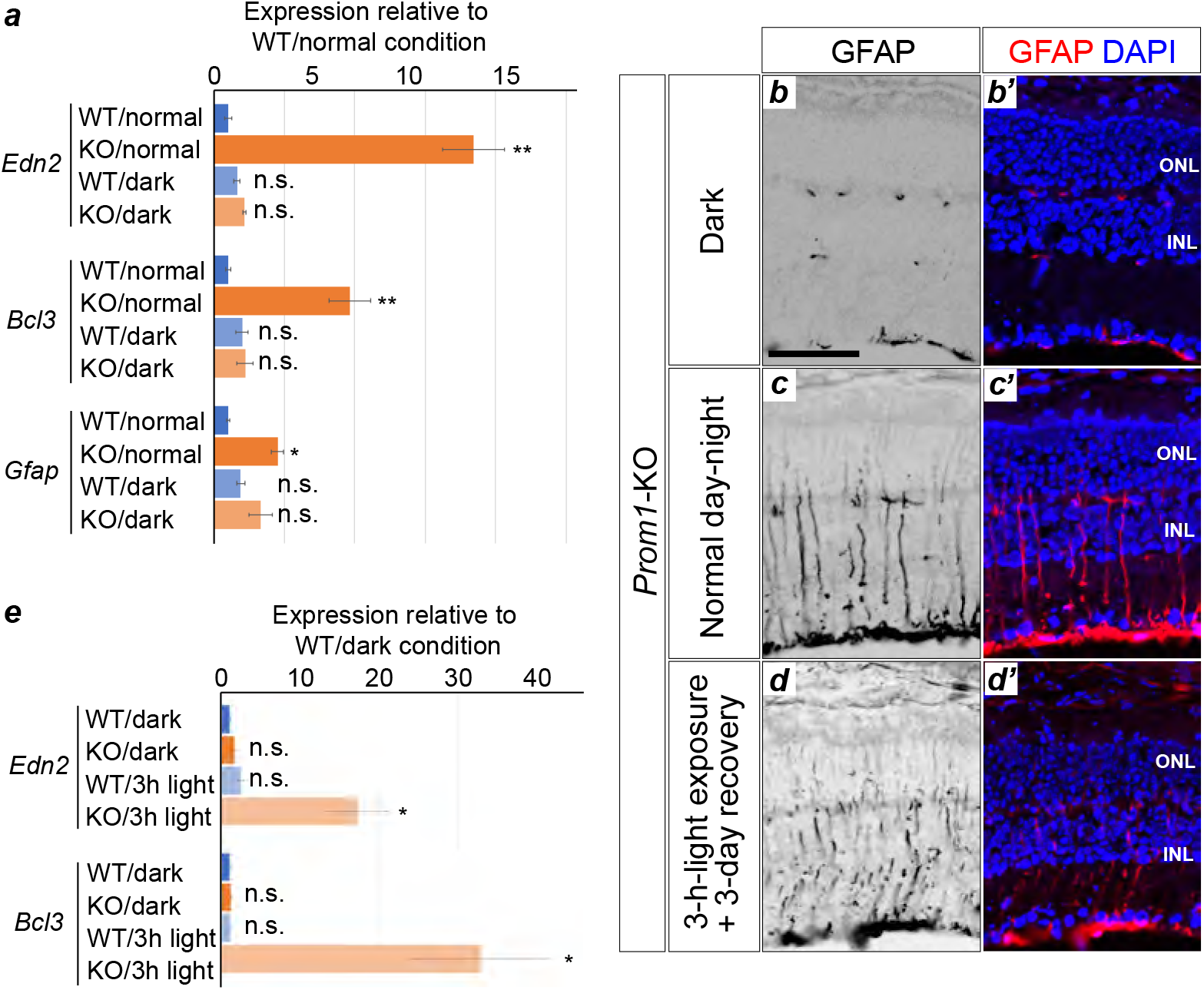
Genes whose expression is increased by Prom1 deficiency are up-regulated by light stimulation. (*a*) RT-qPCR analysis of *Edn2*, *Bcl3*, and *Gfap* expression in the P21 retina of WT or *Prom1*-KO mice that had been reared either under a normal day-night cycle or in the dark. Data are means ± s.e.m. for four retinas for each condition. \**p* < 0.05, \*\**p* < 0.01, n.s., not significant, versus WT/normal (one-way ANOVA followed by Tukey’s post hoc test). (*b* and *c*) Immunofluorescence analysis of GFAP expression in the retina of *Prom1*-KO mice raised as in (*a*). Merged images with DAPI staining are also shown. Scale bar in (b) is 50 μm and applies to all images. Data are representative of four (dark) or seven (normal day-night) retinas. (*d*) RT-qPCR analysis of *Edn2* and *Bcl3* expression in the retina of *Prom1*-KO and WT mice that had been reared in the dark condition for 3 weeks, exposed (or not) to a bright light for 3 h, and then allowed to recover in the dark for 3 days. Data are means ± s.e.m. for five retinas for each condition. \**p* < 0.05, n.s., not significant, versus WT/dark (one-way ANOVA followed by Tukey’s post hoc test). (*e*) Immunofluorescence analysis of GFAP expression in the retina of *Prom1*-KO mice raised in the dark and stimulated with light as in (*d*). Merged images with DAPI staining are also shown. Data are representative of three retinas.

### 3.5. Endothelin receptor antagonists attenuate *Gfap* expression and gliosis in the *Prom1*-KO mouse retina

Endothelin acts at specific receptors [26, 27] to increase both the number of GFAP-positive Müller cells [28] and retinal cell death [29]. Given the elevated expression of *Edn2* and *Gfap* apparent in the retina of *Prom1*-KO mice, we hypothesised that Edn2 might induce aberrant proliferation of glial cells and GFAP expression in association with retinal degeneration in these animals. We therefore examined the possible effects of endothelin receptor antagonists in the mutant mice.

The drugs BQ-123 and BQ-788, which target endothelin receptors A and B, respectively [30], were both injected intraperitoneally into *Prom1*-KO mice at P14, P19, and P24, and the mice were analysed at P28. Whereas GFAP-positive cells were not observed in the retina of WT mice, they were detected in that of *Prom1*-KO mice treated with dimethyl sulphoxide (DMSO) vehicle (figure 5*a* and *b*). However, the number of GFAP-positive cells was markedly reduced in the mutant mice by treatment with BQ-123 and BQ-788 (figure 5*c*). Staining of retinal flat-mount preparations with fluorescently labelled isolectin to detect vascular endothelial cells also revealed fewer retinal vessels in *Prom1*-KO mice than in WT mice and that this difference was attenuated by treatment of the mutant animals with BQ-123 and BQ-788 (figure 5*d*–*g*).

**Figure 5.**
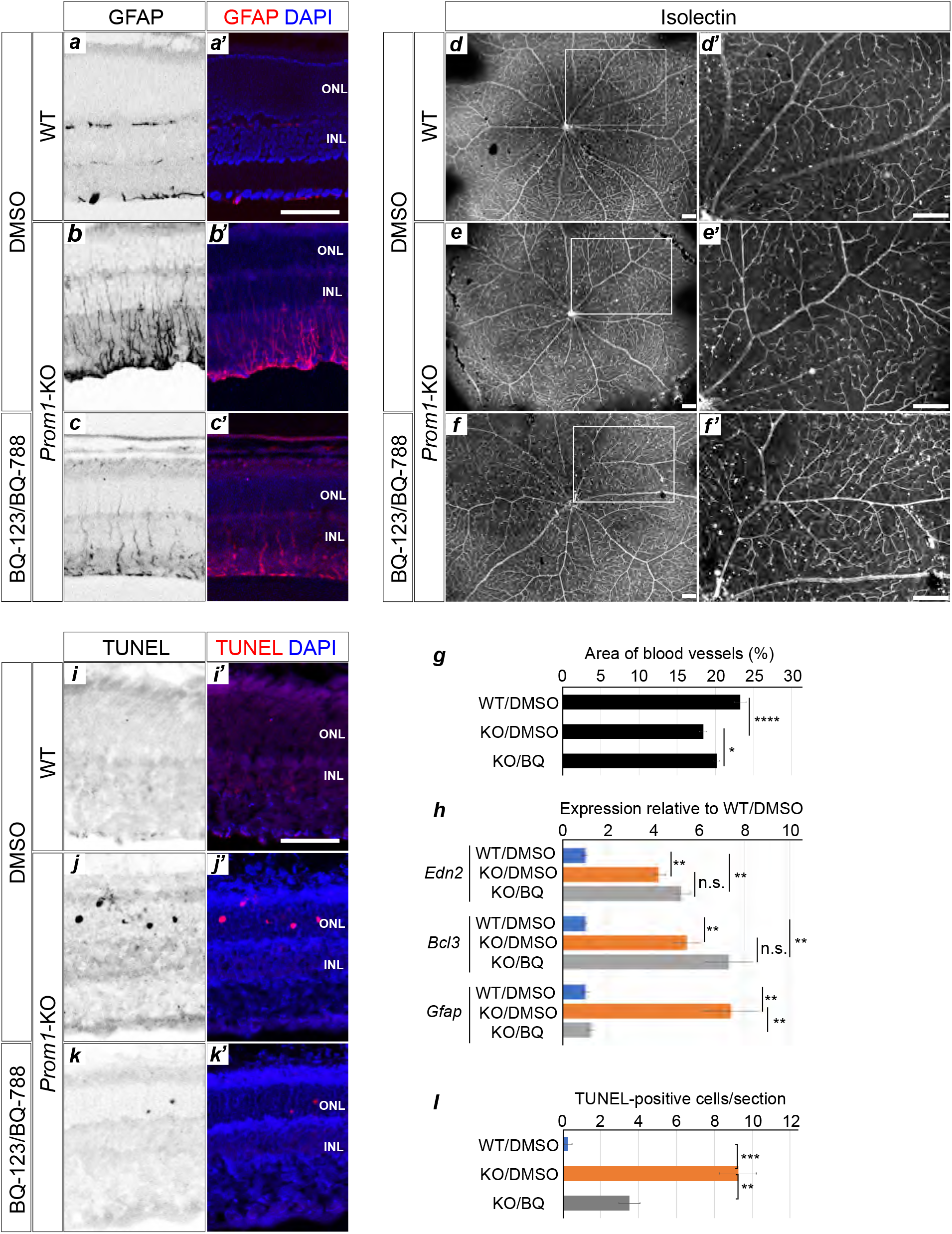
Endothelin receptor antagonists attenuate the increase in the number of GFAP-positive cells and vascular stenosis in the retina of *Prom1*-KO mice. (*a*–*c*) Immunofluorescence analysis of GFAP expression in the retina of WT (*a*) or *Prom1*-KO (*b* and *c*) mice treated with the combination of BQ-123 and BQ-788 (*c*) or with DMSO vehicle (*a* and *b*) at P14, P19, and P24 and analysed at P28. Merged images with DAPI staining are also shown. Scale bar in (a), 50 μm. Data are representative of three retinas per condition. (*d*–*f*) Isolectin staining of the retina of mice as in (*a*) to (*c*). The boxed regions of the left panels are shown at higher magnification in the right panels. Scale bars, 100 μm. (*g*) Area of blood vessels measured in images similar to those in (*d*) to (*f*). Data are means + s.e.m. for X retinas per condition. \**p* < 0.05, \*\*\*\**p* < 0.0001 (one-way ANOVA followed by Tukey’s post hoc test). (*h*) RT-qPCR analysis of *Edn2*, *Bcl3*, and *Gfap* expression in the retina of the treated mice. Data are means ± s.e.m. for three retinas per condition. \*\**p* < 0.01, n.s., not significant (one-way ANOVA followed by Tukey’s post hoc test). (*i*–*k*) TUNEL staining for apoptotic cells in the retina of the treated mice. Merged images with DAPI staining are also shown. Scale bar in (*i*), 50 μm. (*l*) Number of apoptotic cells determined from images similar to those in (*i*) to (*k*). Data are means ± s.e.m. for three retinas per condition. \*\**p* < 0.01, \*\*\**p* < 0.001 (one-way ANOVA followed by Tukey’s post hoc test).

RT-qPCR analysis showed that the expression of *Edn2*, *Bcl3*, and *Gfap* was increased in the retina of *Prom1*-KO mice at P28 compared with that in WT mice. Whereas the expression of *Edn2* and *Bcl3* in the mutant retina was not affected by treatment with BQ-123 and BQ-788, that of *Gfap* was significantly attenuated (figure 5*h*), suggesting that up-regulation of *Gfap* expression in the mutant retina is mediated by endothelin receptor signalling but that that of *Edn2* and *Bcl3* expression is not.

Finally, we examined the effect of BQ-123 and BQ-788 treatment on the number of apoptotic cells in the retina of *Prom1*-KO mice. The TUNEL assay revealed that the marked increase in the number of such cells apparent in the mutant retina at P28 was significantly attenuated by administration of the two drugs (figure 5*l*), suggesting that endothelin receptor signalling contributes to loss of retinal cell homeostasis.

## 4. Discussion

We have here described early manifestations of the retinal degeneration that occurs in *Prom1*-KO mice and identified related genes. We thus detected the aberrant presence of glial cells and the expression of genes associated with the inflammatory response in the mutant retina. Given that the expression of these genes was not activated in the retina of *Prom1*-KO mice maintained in the dark condition, this inflammatory response appears to be dependent on light stimulation. Finally, we found that the deterioration and gliosis characteristic of the mutant retina were ameliorated by the administration of endothelin receptor antagonists.

Although we found that *Prom1* is expressed in the retina from birth, the loss of Prom1 did not substantially affect the expression level of any gene in the retina at P14, suggesting that Prom1 may not play an essential role in the retina prior to light exposure. We previously showed by RT-qPCR analysis that the expression of both *Rdh12* and *Abca4*, two genes that contribute to the visual cycle, was reduced in the retina of *Prom1*-KO mice compared with that of WT mice at P14 [6], suggesting that impairment of the visual cycle might lead to retinal degeneration. Although this result is reproducible as assayed by RT-qPCR (supplementary figure S1), the difference in the expression level of each gene between the two genotypes was associated with a relatively high *p* value in the high-throughput expression analysis performed in the present study (figure 3, supplementary table 2), suggesting this decrease is not critical.

In contrast to the lack of an effect of Prom1 deficiency on the gene expression profile of the retina at P14, we detected many genes, including those related to the inflammatory response, as well as signalling pathways whose activity was altered in the *Prom1*-KO retina at P21. The expression of genes related to phototransduction, for example, was significantly down-regulated in the *Prom1*-KO retina at P21 (figure 3*c*, supplementary table S3), indicating that *Prom1* may be essential for the transcription of such genes or may form a transcriptional network with them. Of note, we found that the expression of causal genes for RP was also down-regulated in the mutant retina at P21.

Of the genes whose expression was up-regulated in the *Prom1*-KO retina at P21, *Edn2* showed the largest fold change. *Edn2* encodes a secretory peptide that plays a role in a wide range of biological processes, including smooth muscle contraction and ovulation [31] as well as development of the enteric nervous system [32]. Its expression is also induced in association with the inflammatory response and promotes glial cell proliferation in the central nervous system [33]. Furthermore, consistent with the perturbation of the retinal vasculature in *Prom1*-KO mice apparent in both the present and a previous [7] study, Edn2 has been found to inhibit retinal vascular development [34]. On the other hand, it was also shown to promote photoreceptor cell survival [35]. These various observations suggest that the role of Edn2 in the photoreceptor degeneration associated with RP and MD is complex.

The expression of *Edn2* has also been shown to be up-regulated in other mouse models of RP [35], including retina-specific *Cdhr1*-KO mice [36], with Prom1 and Cdhr1 having been found to interact with each other [4]. In addition to *Edn2*, the other genes whose expression was affected in the *Prom1*-KO mouse retina overlapped markedly with those affected in the conditional *Cdhr1*-KO mouse retina, suggesting that *Prom1* and *Cdhr1* may function in the same intracellular signalling pathways.

Although we found that the expression of *Edn2* and *Bcl3* in the *Prom1*-KO retina was induced by light stimulation, the mechanism underlying this effect remains unclear. Nevertheless, our study suggests the possibility that an imbalance in intracellular ions caused by the loss of Prom1 (given that Prom1 regulates chloride conductance activated by intracellular calcium uptake [12]) may impair the function of cytoplasmic organelles such as mitochondria and the endoplasmic reticulum, and thereby elicit a stress response. Studies to identify the transcriptional regulatory elements of *Edn2* and the corresponding transcription factors and upstream signalling pathways underlying its photoactivation are warranted.

Gliosis is a response to injury in the central nervous system and is associated with the appearance of GFAP-positive glial cells [26]. It is also a feature of certain neurodegenerative retinal diseases including RP [37], with gliosis in RP having been found to be related to several RP genes. Targeting of gliosis is therefore a potential clinical strategy to delay disease progression and ameliorate associated symptoms. We have now shown that administration of endothelin receptor antagonists attenuated both the appearance of GFAP-positive glial cells and vascular endothelial constriction in the retina of *Prom1*-KO mice. These findings indicate that blockade of endothelin signalling may be an effective clinical strategy for the treatment of gliosis. However, caution is warranted with such an approach for the treatment of RP, given the various functions of endothelins and the consequent potential for adverse systemic effects. Intravitreal injection of endothelin receptor antagonists may help to avoid such side effects. Gene therapy targeting endothelin receptor function is also a potential therapeutic approach for RP. Finally, replacement of dead tissue with functional cells through a regenerative medicine approach may be required for the successful treatment of RP and MD [38].

In conclusion, our results implicating up-regulation of *Edn2* expression in the retinal pathology of *Prom1*-KO mice suggest that localized pharmacological targeting of endothelin receptor signalling warrants further investigation as a clinical intervention for the prevention or treatment of retinal degenerative diseases such as RP and MD.

## Ethics

All animal experiments were approved by the animal welfare and ethics committees of both Yamaguchi University (approval numbers J16021 and U16005 for K.K.) and Nara Institute of Science and Technology (approval numbers 1810 and 311 for N.S.) and were performed in accordance with the relevant guidelines and regulations.

## Data availability

Data are available in the main text/figures and in the Supplementary Information.

## Acknowledgements

We thank Erika Yoshihara, Yukari Mizuno, and Ayaka Kataoka for technical assistance as well as other laboratory members for their support and discussion.

## Author Contributions

KK and NS conceived the project; YK, NS, SW, MS, CY, TO, FH, TY performed experiments; YK, NS, MS, TH, YA analysed the data; All authors joined the discussion; NS, KK, YK wrote the manuscript.

## Supplementary Figure

**Supplementary figure S1.** Expression of *Rdh12* and *Abca4* is down-regulated in the retina of *Prom1*-KO mice at P14.

## Supplementary Tables

**Supplementary table S1.** Primers used for this study.

**Supplementary table S2.** RNA-sequencing analysis of the retina of *Prom1*-KO and WT mice at P14.

**Supplementary table S3.** RNA-sequencing analysis of the retina of *Prom1*-KO and WT mice at P21.

